# Rational design of oxidation-resistant antibodies through local electrostatic modulation

**DOI:** 10.1101/2025.06.29.662139

**Authors:** Sunidhi Lenka, Shrenik Mehta, Nicole Stephens, Franziska Seeger, Wei-Ching Liang, Andrew Watkins, Jonathan Zarzar, Negar Jafari, Rhyan Puno, Caleigh Azumaya, Ryan Kelly, Shuang Wu, Cecilia Chiu, Meredith Hazen, Yan Wu, Flaviyan Jerome Irudayanathan, Sreedhara Alavattam, Bob Kelley, Saeed Izadi

## Abstract

Oxidation is a significant degradation pathway in proteins, particularly therapeutic antibodies, that can impair function, efficacy, and stability. Understanding the sequence and structural factors that drive oxidation susceptibility is critical for incorporating chemical stability into early-stage antibody design. Here, we present a machine learning classifier trained on expert-guided structural features to assess tryptophan (Trp) oxidation risks in the complementarity-determining regions (CDR) of 187 antibodies produced internally at Genentech. The model reveals a strong correlation between local negative electrostatic potential and Trp oxidation susceptibility. A simplified two-parameter model derived from these insights achieves 79% accuracy in classifying oxidation risk, compared to 84% accuracy with the full-feature classifier. Beyond our internal dataset, the two-parameter model successfully predicted oxidation risk for all CDR Trp sites in a blind subset of eight clinical-stage antibodies. In addition, we show that modulating the electrostatic potential around the Trp side-chains through distant charge-altering mutations can significantly reduce oxidation rates. In four out of five re-engineered antibodies, oxidation rates decreased by approximately 50%, while half of these maintained binding affinity. Finally, we demonstrate that this approach can guide multi-property optimization by balancing oxidation resistance and affinity in an anti-CD33 antibody. These results establish a strong link between local electrostatic environments and Trp oxidation susceptibility, and provide a practical framework for designing oxidation resistant biotherapeutics.

## Introduction

Oxidation is a major degradation pathway in proteins with far-reaching implications in physiological functions and pathological conditions. ^1–9^ Oxidative damage is thought to contribute to aging^2^ and the development of diseases such as Alzheimer’s, ^3,4^ ALS, ^5^ and cataracts by increasing protein susceptibility to proteolysis ^6,7^ and reducing enzymatic activity. ^8,9^ In therapeutic antibodies, oxidative damage can compromise structural integrity and function, potentially leading to adverse outcomes including loss of binding affinity, ^10–14^ aggregation, ^13,15,16^ fragmentation, ^17,18^ color change of drug substance ^19,20^ and altered immunogenicity. ^21^ Understanding the determinants and underlying mechanisms of oxidation and the associated risks are critical for assessing its role in disease progression and ensuring the long-term chemical and physical stability of therapeutic antibodies. ^22–24^

Among the amino acids prone to oxidation, methionine (Met) and tryptophan (Trp) stand out as particularly susceptible in antibody therapeutics. ^13,23,25,26^ Met oxidation has been extensively studied, with its susceptibility closely tied to solvent accessibility. ^27–29^ In contrast, Trp oxidation is less well understood yet arguably more consequential in therapeutic contexts ^12,14,25^ for two key reasons. First, Trp oxidation involves multiple reaction pathways and produces a variety of oxidative products, including oxindolealanine, hydroxytryptophan, kynurenine, and N-formylkynurenine (Figure 1a). This complexity makes predicting and characterizing Trp oxidation far more challenging. ^14,30^ Second, Trp plays a critical role in protein-protein and hydrophobic core interactions, ^31–33^ owing to its unique structural properties. Trp’s side chain features a highly accessible hydrophobic surface, a strong potential for cation–*π* interactions, and an indole N–H group capable of hydrogen bonding. In fact, the unique properties makes Trp more prevelant,^34^ and highly surface exposed, ^25,35^ in the complementarity-determining regions (CDRs) of antibodies compared to other proteins.

**Figure 1:**
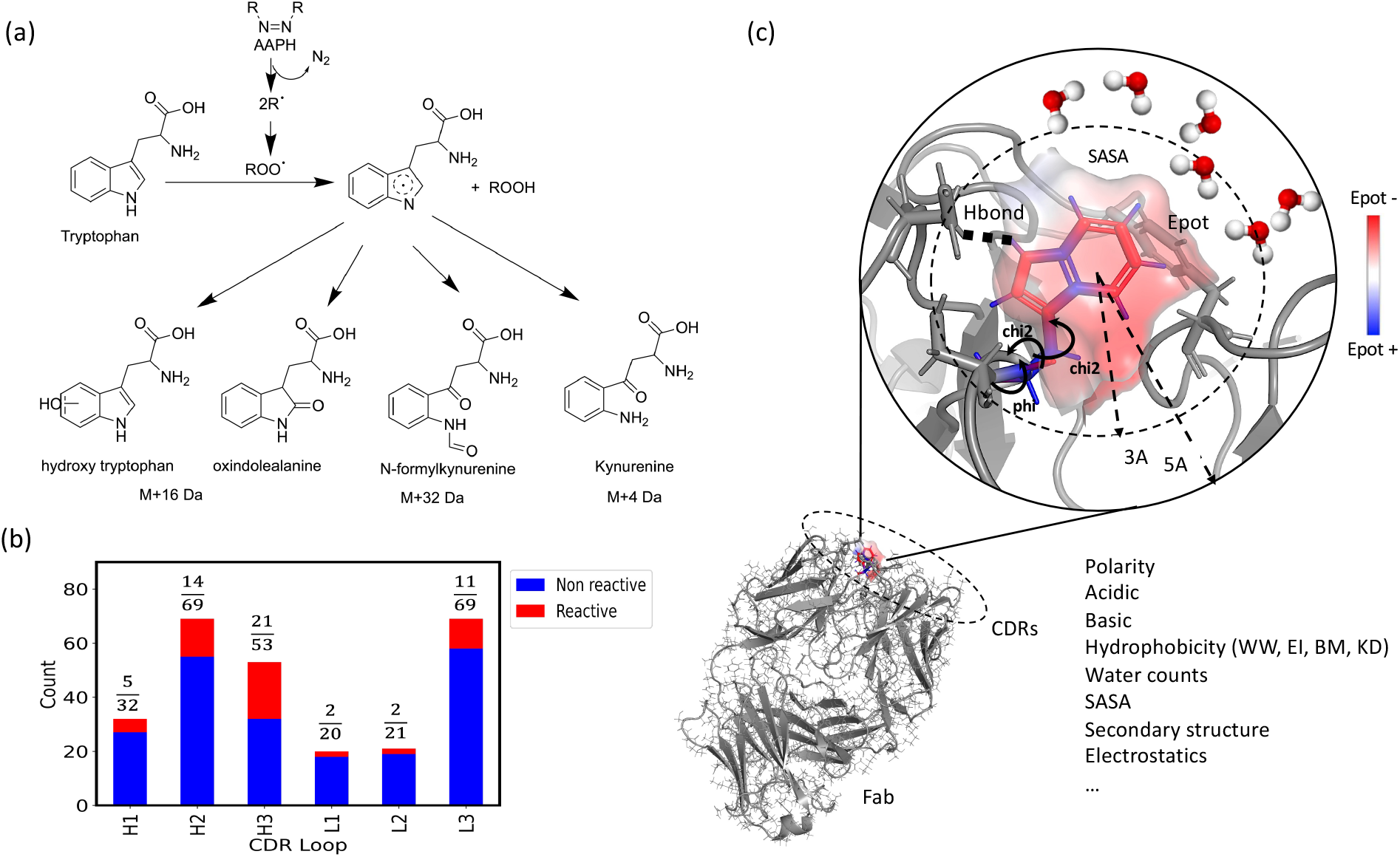
(a) Representative oxidation products of Trp under AAPH-mediated oxidative stress. The AAPH reagent generates ROO^*·*^ radicals that react with Trp to form a Trp radical intermediate, which can further oxidize to produce 4-, 5-, 6-, and 7-hydroxytryptophan (+16 Da), oxindolealanine (+16 Da), N-formylkynurenine (+32 Da), and kynurenine (+4 Da). (b) Frequency of Trp oxidation observed in the dataset as a function of CDR loop location under AAPH stress conditions. A 35% oxidation threshold was applied to classify Trp residues as oxidized (red) or unoxidized (blue). (c) Overview of structural and surface descriptors used in the machine learning model to classify oxidation risk for CDR Trp residues.

Computational models ranging from quantum mechanical calculations ^36^ to molecular dynamics simulations ^29,37^ and machine learning approaches ^27,38^ have been used to probe the mechanisms of protein oxidation and to develop predictive models. Among the key factors influencing oxidation susceptibility, the solvent-accessible surface area (SASA) of Trp and Met has been consistently identified as a critical determinant.^36–41^ Additionally, structural parameters such as dihedral angles (*ϕ* and *ψ*), ^38^ torsional properties, ^38^ loop flexibility, ^36^ and proximity to aromatic residues ^38,42,43^ have also been investigated, albeit to a lesser extent.

Design strategies to mitigate or prevent Trp oxidation are as critical as predictive efforts; however, they remain relatively limited. One common approach involves substituting oxidation-prone Trp residues with other aromatic amino acids, such as phenylalanine or tyrosine. ^12,44^ Nevertheless, due to the distinct physicochemical properties of Trp, its removal is often impractical, as it can adversely affect antigen binding ^45,46^ and specificity. ^47^ Therefore, rational engineering strategies that preserve Trp within the binding site while mitigating its susceptibility to oxidation are highly desirable.

In this study, we leverage a large dataset of antibody Trp oxidation profiles to identify new structural factors that influence oxidation susceptibility. Our findings highlight the critical role of local electrostatics and demonstrate how this feature can be exploited to rationally design oxidation resistant antibodies. Through point mutations introduced distally to the binding site Trp residues, we show that oxidative stability can be significantly improved while maintaining binding affinity. We further present a multi-parameter optimization framework that systematically enhances oxidation resistance while minimizing trade-offs typically associated with Trp substitution.

## Results

### Oxidation susceptibility of CDR tryptophan residues

We analyzed a dataset of 187 monoclonal antibodies (mabs) produced internally at Genentech, which contained 264 Trp residues within their CDRs. These antibodies were exposed to oxidative stress using AAPH as a radical generator and the level of oxidation was measured as a percentage of modified Trp residues post stress. Using a historically established threshold of 35% modifications, ^48^ we categorized Trp residues as either susceptible (high risk, n = 55) or resistant (low risk, n = 209) to oxidation.

Figure 1b shows the distribution of oxidation risk across different antibody CDR loops. Overall, 20% of all Trp residues within the CDRs were identified as susceptible to oxidation. Oxidation risk varied significantly among different CDR loops. Approximately 40% of Trp sites within the H3 loops underwent oxidation, compared to 20% within the H2 loops, 16% within the L3 and H1 loops, and 10% in the L1 and L2 loops. Given the critical role of the H3 loop for antigen binding, ^49–51^ the high frequency of Trp oxidation in this loop is particularly noteworthy, highlighting the associated impact on overall stability and function of antibodies.

### Structure-based feature engineering, selection, and interpretation

Feature selection was informed by mechanistic insights from prior studies on protein oxidation and stability, ^36–43^ as well as hypotheses specific to Trp oxidation. We incorporated a range of commonly studied structural descriptors, including SASA, backbone dihedral angles, loop flexibility, and the types of neighboring amino acids, features previously shown to influence oxidation susceptibility. ^36–41^

In addition to these established features, we introduced new descriptors relevant to physicochemical stability, including residue-level electrostatic potential (Epot) ^52^ and residue-level spatial aggregation propensity (SAP) scores, ^53^ computed using various hydrophobicity scales. Epot represents the electrostatic potential experienced by each residue in the solvated antibody, calculated via the Poisson–Boltzmann equation. ^54^ SAP quantifies the degree of local clustering of hydrophobic residues on the molecular surface. ^53^ A residue-level SAP determines the hydrophobicity of neighboring residues.

Altogether, we compiled a total of 50 structural features capturing key biophysical and spatial properties, including solvent exposure (e.g., SASA, SASA loop, n loop water 5), flexibility (e.g., loop rmsd, res rmsd), local hydrophobicity (e.g., SAP scores), backbone geometry (e.g., *ψ, ϕ*, and *χ* angles of neighboring residues), local electrostatics (e.g., Epot), and interactions with surrounding polar, nonpolar, and aromatic residues (Figure 1c). All features were averaged over molecular dynamics trajectories, starting from the predicted structures of the Fab domains (see Methods for details). A full list of features is provided in the Supporting Information (Table S1).

To explore inter-feature relationships, we generated a dendrogram (Figure 2a), which revealed eleven distinct feature clusters. These clusters could be broadly categorized into six biophysical groups: solvent accessibility, local electrostatics, local hydrophobic residue clustering, flexibility, backbone geometry, and side-chain dihedral angles.

**Figure 2:**
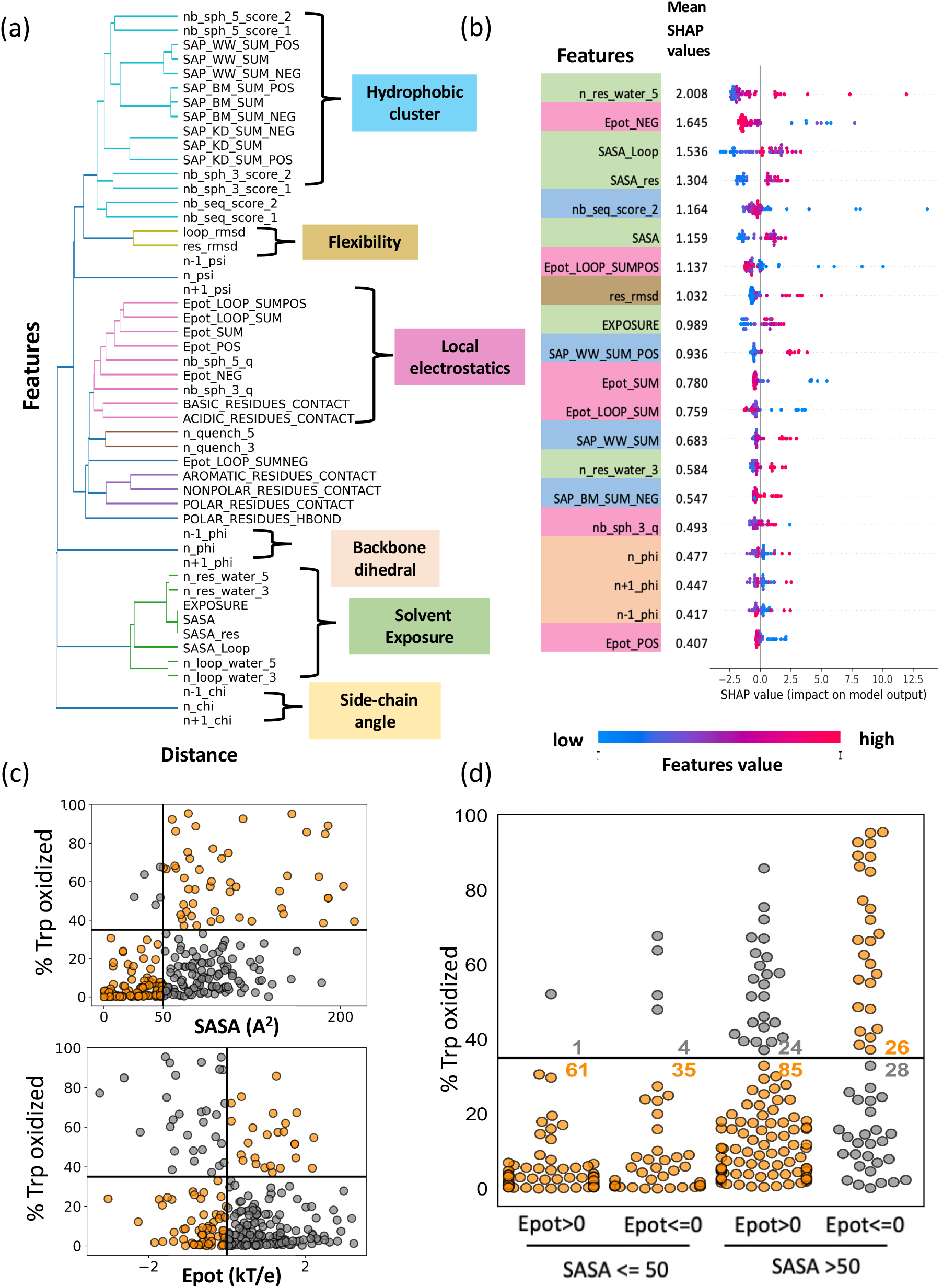
a) Hierarchical clustering of structural and environmental descriptors from Fab domain models revealed 11 distinct feature clusters, grouped into six broader biophysical categories. b) Feature importance rankings derived from Random Forest and SHAP analysis highlight solvent accessibility (SASA) and local electrostatic potential (Epot) as the top two predictors of Trp oxidation susceptibility. Features are colored based on the biophysical categories in panel a. c) Correlation plots showing the relationship between Trp oxidation rates and two key descriptors, SASA and Epot. d) Swarm plots showing the distribution of Trp oxidation rates across different classes of SASA and Epot values. Trp residues that are both highly exposed (SASA *>* 50 Å^2^) and located in a negatively charged environment (Epot *<* 0 kT/e) exhibit the highest oxidation rates, highlighting the combined effect of solvent accessibility and local electrostatics.

Using Random Forest (RF) feature importance analysis and SHAP (SHapley Additive exPlanations) values (Figure 2b), we identified key structural and environmental features associated with Trp oxidation susceptibility. As expected, solvent accessibility, captured by features such as n res water 5, SASA Loop, and SASA res, emerged as the most influential factor. This is consistent with prior findings, as Trp residues buried within the protein structure are shielded from AAPH-derived radicals and therefore less prone to oxidation.

Notably, electrostatic potential was the second most important feature category, despite being relatively unexplored in the context of Trp oxidation. Features such as Epot NEG, Epot LOOP SUMPOS, Epot SUM, and Epot LOOP SUM showed strong predictive power. SHAP analysis revealed that more negative electrostatic potentials around the Trp side-chain (e.g., high Epot NEG) are associated with increased oxidation risk, whereas positive potentials (e.g., Epot POS) appear protected.

Other features also contributed meaningfully, though to a lesser extent. Local clustering of hydropho-chain dihedral angles.

Using Random Forest (RF) feature importance analysis and SHAP (SHapley Additive exPlanations) values (Figure 2b), we identified key structural and environmental features associated with Trp oxidation susceptibility. As expected, solvent accessibility, captured by features such as n res water 5, SASA Loop, and SASA res, emerged as the most influential factor. This is consistent with prior findings, as Trp residues buried within the protein structure are shielded from AAPH-derived radicals and therefore less prone to oxidation.

Notably, electrostatic potential was the second most important feature category, despite being relatively unexplored in the context of Trp oxidation. Features such as Epot NEG, Epot LOOP SUMPOS, Epot SUM, and Epot LOOP SUM showed strong predictive power. SHAP analysis revealed that more negative electrostatic potentials around the Trp side-chain (e.g., high Epot NEG) are associated with increased oxidation risk, whereas positive potentials (e.g., Epot POS) appear protected.

Other features also contributed meaningfully, though to a lesser extent. Local clustering of hydrophobic and aromatic residues, captured by metrics such as nb seq score 2 and SAP scores, was associated with elevated oxidation susceptibility. Additionally, greater local flexibility, as indicated by higher res rmsd values, correlated with increased oxidation risk. These observations are in line with previous studies highlighting the role of loop dynamics^36^ and neighboring aromatic interactions ^38,42,43^ in modulating oxidation behavior.

### Assessing the role of key biophysical descriptors in Trp oxidation risk

To better understand the individual predictive contributions of key features and to reduce model complexity, we evaluated a simplified two-parameter model based on the top-ranked descriptors: solvent accessibility (SASA) and local electrostatics (Epot). We compared the performance of this minimal model against that of full-feature RF and Gradient Boosting (GB) classifiers, which were trained using all 50 features and evaluated via 10-fold cross-validation (Table 1). The high-feature models achieved an accuracy of approximately 84%, with high specificity (96% for RF and 94% for GB), but limited sensitivity (40% and 45%, respectively), reflecting a tendency to underpredict oxidation-prone residues (i.e., a high false negative rate).

**Table 1:**
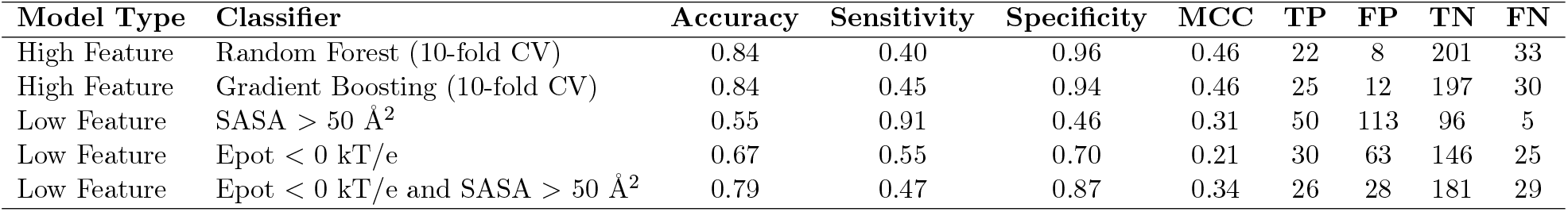
Performance comparison of machine learning models and rule-based classifiers for predicting Trp oxidation.

| Model Type | Classifier | Accuracy | Sensitivity | Specificity | MCC | TP | FP | TN | FN |
| --- | --- | --- | --- | --- | --- | --- | --- | --- | --- |
| High Feature | Random Forest (10-fold CV) | 0.84 | 0.40 | 0.96 | 0.46 | 22 | 8 | 201 | 33 |
| High Feature | Gradient Boosting (10-fold CV) | 0.84 | 0.45 | 0.94 | 0.46 | 25 | 12 | 197 | 30 |
| Low Feature | SASA > 50 Å<sup>2</sup> | 0.55 | 0.91 | 0.46 | 0.31 | 50 | 113 | 96 | 5 |
| Low Feature | Epot < 0 kT/e | 0.67 | 0.55 | 0.70 | 0.21 | 30 | 63 | 146 | 25 |
| Low Feature | Epot < 0 kT/e and SASA > 50 Å<sup>2</sup> | 0.79 | 0.47 | 0.87 | 0.34 | 26 | 28 | 181 | 29 |

We next examined the predictive utility of the top features individually (Figure 2c and Table 1). A correlation analysis between SASA and observed oxidation frequency revealed a positive relationship, indicating that Trp residues with greater solvent exposure are more likely to undergo oxidation. To investigate this further, we categorized Trp residues as either buried or exposed based on their SASA values. Guided by the correlation plot, we empirically defined buried Trp residues as those with SASA *<*= 50 Å^2^, a threshold consistent with the relative solvent accessibility (RSA) cutoffs of *<*20–25% reported in prior studies. ^55^

Consistent with expectations, buried Trp residues exhibited a strong protective effect: 96 out of 101 were correctly classified as non-oxidized (true negatives). This aligns with the mechanistic understanding that burial limits access to AAPH-derived radicals. However, exposure alone was not sufficient to predict oxidation. Among the 163 exposed Trp sites (SASA *>* 50 Å^2^), only 50 were oxidized, corresponding to a specificity of just 46%. This indicates that while solvent exposure is a necessary condition, additional factors also play a critical role in determining oxidation susceptibility.

The second most important feature, electrostatic potential (Epot), exhibited a negative correlation with oxidation rate. Specifically, approximately 85% of Trp residues situated in a positively charged environment (Epot *>* 0) were correctly classified as non-reactive (146 out of 171 cases; Figure 2c, Table 1). As a single-feature classifier, Epot outperformed SASA, achieving higher overall accuracy (67% vs. 55%) and greater specificity. This suggests that a locally positive electrostatic environment may offer a protective effect against oxidation. However, Epot showed lower sensitivity than SASA, indicating it is less effective at detecting oxidation-prone residues.

Combining SASA and Epot substantially improved predictive performance over either feature alone. As shown in Figure 2d, 85 out of 109 (79%) exposed Trp residues in a positively charged environment fell below the oxidation rate threshold, compared to only 28 out of 54 (52%) of exposed sites in a negatively charged environment. The resulting two-feature classification model achieved an overall accuracy of 79%, closely approaching the performance of the full-feature RF and GB models (84%), while using only a fraction of the input complexity (Table 1). These results underscore the dominant role of solvent accessibility and local electrostatics in governing Trp oxidation susceptibility and demonstrate that much of the predictive power of the high-dimensional models can be captured using these two interpretable biophysical descriptors.

### Generalizability of the low-feature model: blind prediction on clinical antibodies

To assess the generalizability of our two-feature model beyond the Genentech internal dataset, we conducted a blind prediction study on a set of clinical-stage antibodies. We curated a panel of 629 clinical-stage IgG1 molecules from a 2022 snapshot of TheraSAbDab, ^56^ extracting variable domain sequences. For each antibody, static structure models of the Fab domains were generated using DeepAb ^57^ (see Methods). SASA and electrostatic potential (Epot), the top predictive features identified earlier, were calculated for all Trp residues located within the CDR loops. Figure 3a shows a two-dimensional scatter plot of all CDR Trp residues across the dataset, based on their SASA and Epot values.

**Figure 3:**
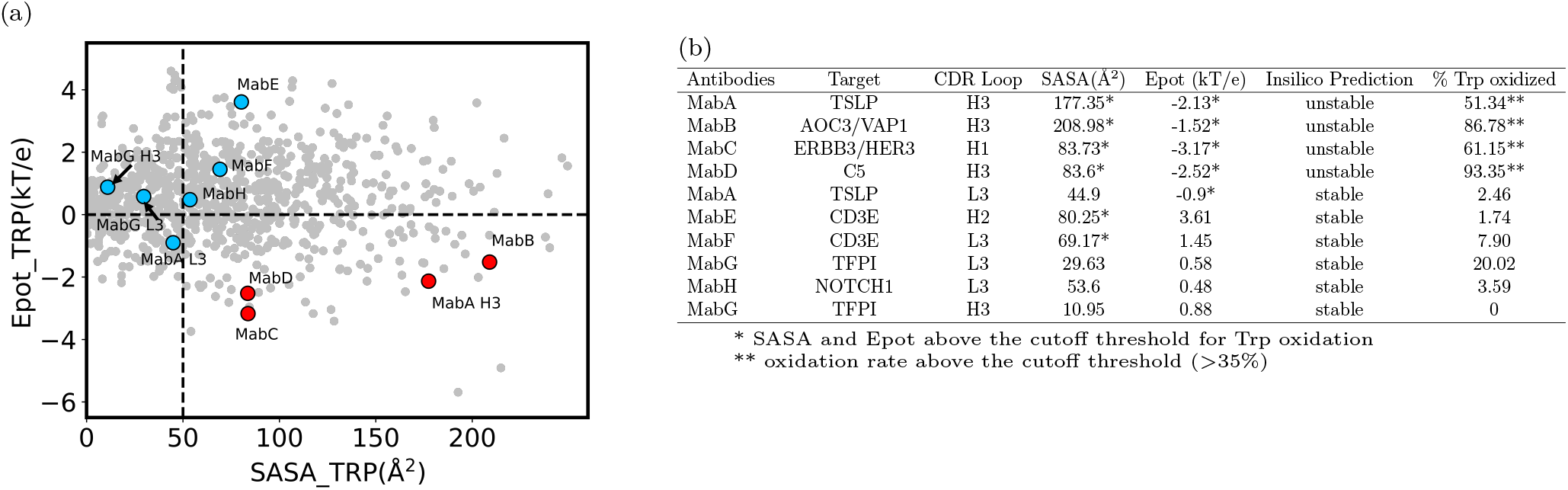
a) Distribution of clinical-stage antibodies based on solvent-accessible surface area (SASA) and electrostatic potential (Epot) of CDR Trp residues. Ten cases from eight clinical IgG1 antibodies were selected for AAPH-mediated Trp oxidation stress studies. Red points indicate Trp residues with high oxidation (*>*35%), while blue points represent low oxidation (*<*35%) observed after AAPH stress. b) Summary of SASA, Epot descriptors, and experimentally measured oxidation rates for the selected clinical antibodies. SASA and Epot values were averaged over molecular dynamics (MD) trajectories. In silico predictions of oxidation susceptibility showed strong agreement with experimental outcomes.

To guide selection for experimental validation, we divided the plot into four quadrants using SASA = 50 Å^2^ and Epot = 0kT/e as empirical thresholds. From each quadrant, representative antibodies were randomly selected, with particular emphasis on candidates predicted to be at high oxidation risk, i.e., those with Trp residues in highly exposed (SASA *>* 50Å^2^) and negatively charged (Epot *<* 0kT/e) environments, marked as red points in Figure 3a. The sequences for the 8 selected clinical antibodies are provided in the SI (Table S4).

A total of eight antibodies, all sharing the same IgG1 constant framework but with distinct variable regions, were expressed and subjected to AAPH oxidative stress under standardized conditions. Trp oxidation levels were then experimentally quantified and compared against our a priori in-silico predictions.

Figure 3b summarizes the SASA and Epot values, predicted oxidation risks, and experimentally measured Trp oxidation rates for ten sites across these eight antibodies. Notably, our model correctly classified a ll t en T rp r esidues. I n p articular, t he four residues with high SASA and negative Epot values exhibited substantial oxidation (51.34% to 93.35%), in line with our predictions. These included MabA (H3 loop), MabB, MabC, and MabD, all of which contained CDR Trp residues predicted to be unstable due to their high exposure and unfavorable electrostatic environment.

Conversely, Trp residues in positively charged or buried environments were oxidation-resistant. For example, MabE and MabF displayed high SASA but were located in regions with positive Epot values, and both showed minimal oxidation. Similarly, MabA contained a second Trp residue with borderline SASA (44.9 Å^2^) and mildly negative Epot, which also remained stable after AAPH treatment.

These results demonstrate that a simple two-feature model, using only solvent accessibility and local electrostatics, can robustly predict Trp oxidation outcomes in external datasets. The successful blind prediction of oxidation susceptibility across clinical-stage antibodies underscores both the generalizability of the model and the mechanistic relevance of the selected features.

### Electrostatic modulation via distal point mutations reduces Trp oxidation risk

Given the strong influence of local electrostatics on predicting Trp oxidation risk, we investigated whether modulating the strength and sign of the electrostatic potential around susceptible Trp residues through point mutations would affect oxidation rates. To test this, we analyzed a subset of five Genentech internal antibodies with CDR Trp residues within a negative electrostatic potential and oxidation rate *>*35%. To mitigate the oxidation risk, we introduced point mutations by substituting nearby acidic residues with basic residues. Specifically, single, double and triple D → R/K mutations were applied to shift the local electrostatic potential around the Trp side-chain from negative to positive. Note that our designs focused exclusively on removing or mitigating the oxidation risk, without necessarily optimizing or considering the impact on binding affinity.

Table 2 and Figure 4 summarize the calculated electrostatic descriptors (Epot), solvent accessibility (SASA), and oxidation rates for the original antibodies and their engineered variants. In Mab1, W91 in the L3 loop has highly negative electrostatic potential (−1.55 kT/e) and a SASA of 58.38 Å^2^, resulting in a high oxidation rate of 70.6%. To counteract this, we identified three Asp residues in the H3 loop and mutated them to Arg (D97R, D99R & D101R). This triple mutation shifted the side-chain electrostatic potential to a mildly positive value (0.69 kT/e) and reduced SASA (28.5 Å^2^). The impact on SASA was likely due to increased packing and conformational adjustments. Through these mutations, the oxidation rate decreased by 46%, lowering to 38.1% in the triple variant Mab1.RRR.

**Table 2:** Summary of mutations, electrostatic potential, SASA, oxidation rates for CDR Trp residues in internal antibodies. The change in antigen binding kinetics (measured by SPR at pH 7.4) upon point mutations relative to the wild type is also reported.

| Molecule | Trp residue (Kabat) | AA mutations (Kabat) | Epot of Trp (kT/e) | SASA of Trp ($\text{\AA}^2$) | Oxidation rate | Relative KD |
| --- | --- | --- | --- | --- | --- | --- |
| Mab1 | W91(L3) | H3: D97D99D101 (WT) | -1.55 | 58.38 | 70.6% | 1 |
| Mab1.RRR | W91(L3) | H3: R97R99R101 | 0.69 | 28.50 | 38.1% | 60 fold decrease |
| Mab2 | W58(H2) | H3: R100aD100cD100dD101 (WT) | -1.52 | 80.53 | 45.6% | 1* |
| Mab2.RRR | W58(H2) | H3:R100aR100cR100dR101 | 1.65 | 66.04 | 28.3% | NB |
| Mab2.W | W100a(H3) | H3: W100aD100cD100dD101 | -2.22 | 220.81 | 96.8% | – |
| Mab3 | W100b(H3) | H3:D98(WT) | -1.61 | 197.14 | 56.2% | 1 |
| Mab3.R | W100b(H3) | H3:R98 | 1.23 | 192.50 | 53.3% | 1.4 fold decrease |
| Mab4 | W100a(H3) | H3: D95D100D101(WT) | -1.24 | 64.20 | 86.2% | 1 |
| Mab4.K | W100a(H3) | H3: D95K100D101 | -0.66 | 60.58 | 41.7% | 2.2 fold decrease |
| Mab4.RR | W100a(H3) | H3: R95D100R101 | 1.17 | 107.85 | 1.3% | NB |
| Mab4.KR | W100a(H3) | H3: D95K100R101 | -0.24 | 112.51 | 16.6% | NB |
| Mab5 | W33(H1) | H1: D31Y32(WT) | 0.29 | 81.33 | 40.7% | 1 |
| Mab5.A | W33(H1) | H1: A31Y32 | 1.21 | 77.76 | 34.7% | 3.4 fold decrease |
| Mab5.AA | W33(H1) | H1: A31A32 | 1.31 | 75.02 | 18.7% | 2 fold decrease |

NB:No binding observed
\*KD could not be accurately determined as off rate was reaching the limit of the instrument

**Figure 4:**
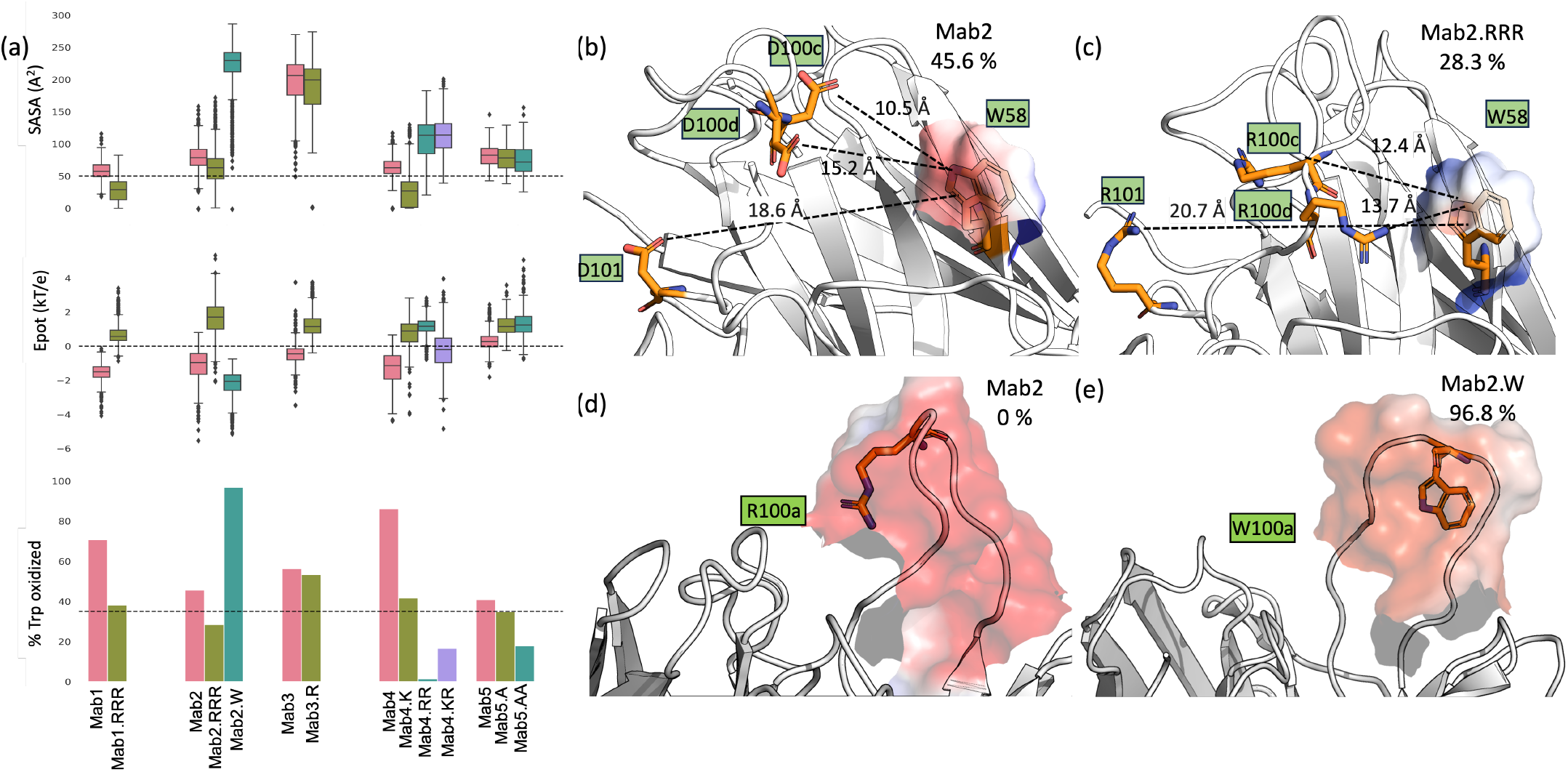
a) SASA, electrostatic potential (Epot), and Trp oxidation rates under AAPH stress for antibody case studies and their designed variants. Oxidation decreases in variants where Trp resides in a more positive electrostatic environment. b) Visualization of the negative electrostatic potential surrounding Trp in Mab2. c) Triple Asp→Arg mutations at *>*10 Å from Trp in Mab2.RRR increase local Epot and reduce oxidation by 37.9%. d) Concentrated negative electrostatic potential in the H3 loop of Mab2. e) Introduction of Trp100a into the negatively charged H3 loop of Mab2 leads to 96.8% oxidation under AAPH stress.

In Mab2, a similar approach was taken to reduce the oxidation rate of W58 in the H2 loop, initially at 45.6%. Three D → R mutations in the H3 loop were introduced (D100cR, D100dR, D101R), shifting the local electrostatic potential significantly from -1.52 kT/e to 1.65 kT/e. Unlike in Mab1, the SASA for this variant remained high (66.04 Å^2^ compared to 80.53 Å^2^ for the wild type), and the mutations were positioned at distances ranging from 10.5 Å to 18.6 Å from the Trp site (Figure 4b). Despite the high SASA in the variant Mab2.RRR, the oxidation rate decreased from 45.6% to 28.3% (below the cutoff).

For Mab3, we introduced a single D98R mutation, which significantly altered the electrostatic potential but had minimal impact on SASA. The reduction in the oxidation rate of W100b in H3 was marginal, with only a 2% decrease, suggesting that altering electrostatics alone was insufficient in this case. We speculate that the extremely high SASA (close to RSA of 90% or 197 Å^2^) may be the dominant factor, over-shadowing the effects of changes in electrostatic potential and limiting the impact on oxidation.

In contrast, W100a in Mab4’s H3 loop exhibited a pronounced response to electrostatic modulation. A single D100K mutation (Mab4.K) reduced the oxidation rate by more than 50%, from 86.2% to 41.7%. A double mutation (D95R and D100R) nearly eliminated the oxidation risk, lowering the rate to just 1.3%. Notably, this double mutation (Mab4.RR) also increased SASA by 78% compared to the wild type (RSA = 49% or SASA = 107.85 Å^2^), suggesting that for intermediate RSA values, electrostatic potential becomes the dominant factor in oxidation susceptibility. Interestingly, the Mab4.KR double mutation shifted the side-chain Epot to a small negative values (−0.24 kT/e), compared to Mab 4 Epot. This modification was still highly effective in lowering the oxidation rate by approximately 80%, reducing it to 16.6% though the oxidation is higher than Mab4.RR where the side chain Epot is positive.

In Mab5, a single D31A mutation reduced the oxidation rate of W33 (H1) from slightly above the threshold (40.7%) to just below it (34.7%). We observed that the presence of Y32 in the n-1 position might be shielding the effect of the D31A mutation on W33. To further investigate this, we introduced a Y32A mutation in combination with D31A, which resulted in an increase in the positive potential of the W33 site. The double D31A and Y32A mutation further reduced the oxidation rate by 54%, bringing it down to 18.7%. This combination underscores the synergistic effects of aromatic clustering (third important feature in our ML model) and charge modulation in mitigating oxidation risk. Overall, these experimental validations confirm that targeted distant electrostatic mutations can effectively reduce Trp oxidation.

### Engineering an oxidation-prone Trp through electrostatic contextual insertion

In addition to engineering strategies aimed at mitigating Trp oxidation, we sought to test whether we could intentionally induce oxidation susceptibility by introducing a Trp residue into an unfavorable local electrostatic environment. Specifically, we aimed to validate whether placing a Trp residue within a negatively charged, solvent-exposed surface patch would be sufficient to trigger high oxidation rates, independent of native sequence or structural context.

To test this, we identified residue R100a in the H3 loop of Mab2, a site located at the center of a strongly negative electrostatic potential (Figure 4d). We chose Arg as the starting residue due to its structural similarity to Trp in terms of bulky size and solvent accessibility, as well as its inherent positive charge, making it an ideal candidate to assess the effect of charge reversal.

We introduced a single point mutation, R100aW, generating the variant Mab2.W (Figure 4e and Table 2). The resulting Trp residue (W100a) had a high solvent-accessible surface area (SASA = 220.81 Å^2^) and a strongly negative electrostatic potential (Epot = –2.22 kT/e). Experimental evaluation of this variant under AAPH-induced oxidative stress revealed an extremely high oxidation rate of 96.8%, confirming that the engineered Trp residue was indeed highly susceptible to oxidation.

This finding provides compelling, mechanistic evidence that solvent accessibility and local electrostatics are sufficient, independent of neighboring sequence motifs or structural elements, to drive Trp oxidation.

### Binding affinity assessment of oxidation-reducing variants

Given the success of electrostatic point mutations in reducing Trp oxidation, we next evaluated whether these modifications impacted antigen binding. Importantly, the original designs were made without any consideration of binding affinity; the sole objective was to mitigate oxidation risk by modulating local electrostatics.

Despite this, binding was retained in 4 out of the 8 engineered variants (50%), as summarized in Table 2. Notably, Mab4.K and the Mab5 variants (Mab5.A and Mab5.AA) demonstrated substantial reductions in oxidation while maintaining strong binding to their respective targets. Mab3.R also retained binding affinity, although it showed only marginal reduction in oxidation risk. These results suggest that certain electrostatic mutations, even when applied to CDR-adjacent or framework regions, can be tolerated without compromising antigen recognition.

### Dual optimization of Trp stability and binding affinity in an anti-CD33 antibody

Given the promising results from oxidation-reducing variants that retained binding, we next explored whether it was possible to simultaneously optimize both Trp oxidation resistance and antigen-binding affinity. These efforts focused on a clinically relevant antibody targeting CD33, where a critical tryptophan residue (W96) in the CDR H3 loop was identified as highly oxidation-prone and essential for binding, see Figure 5.

**Figure 5:**
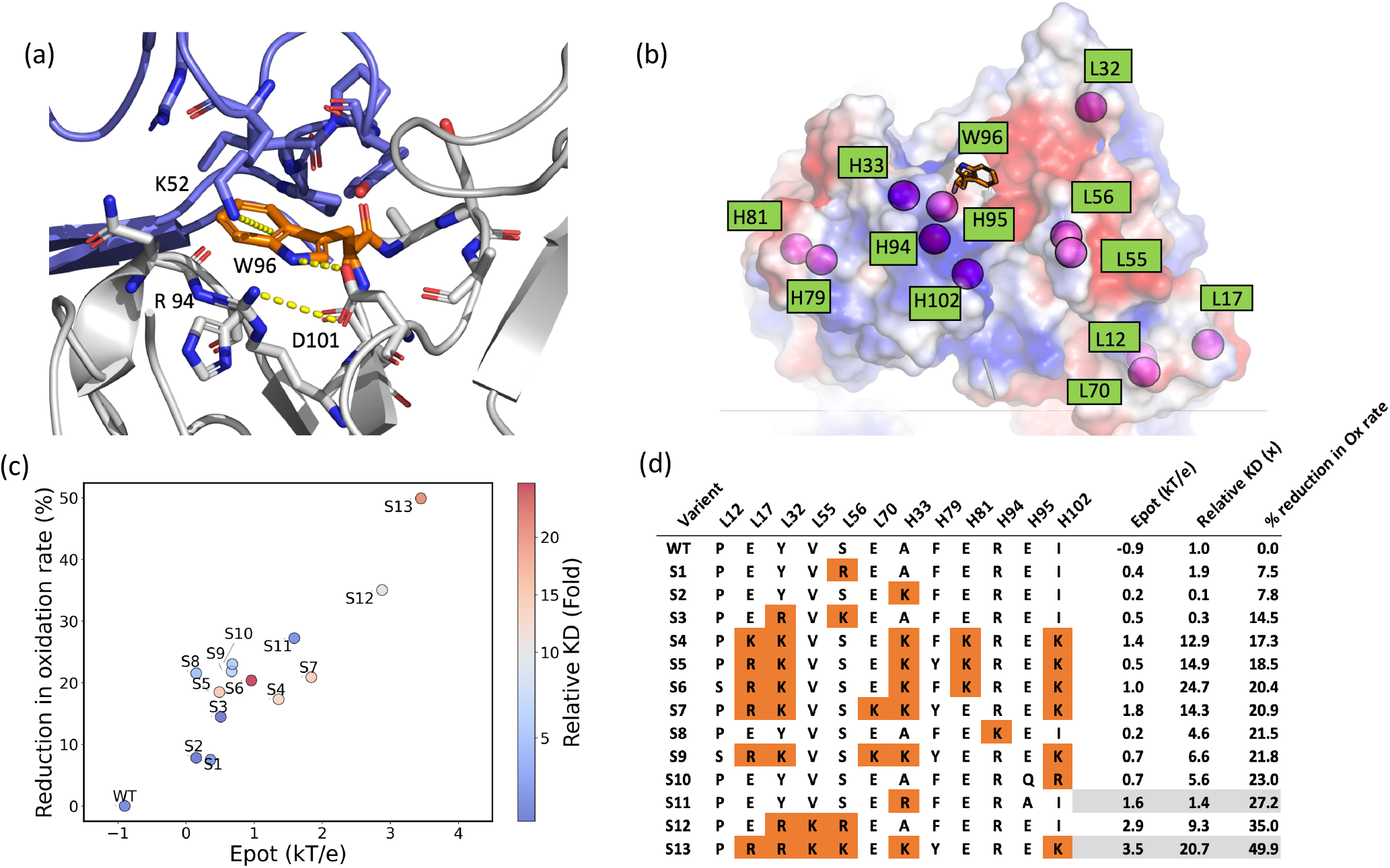
a) Cryo-EM structure of 15G15.3 Fab (gray) bound to CD33 (violet), showing Trp96 forming key interactions with Lys52 (CD33) and Asp101 (Fab). Trp96 oxidizes at 97% under AAPH stress; W96F mutation abolishes binding (*>*1000-fold loss). b) Electrostatic potential of the lead candidate, with 12 mutated residues shown as pink spheres. c) Scatter plot of Trp oxidation vs. Epot for the lead and 13 variants; point color reflects relative KD. d) Summary of 13 engineered variants. S11 and S13 show improved oxidation resistance with preserved binding.

W96 is situated at the VH–VL interface and plays a central role in antigen recognition. Structural analysis using a Fab–CD33 antigen CryoEM complex (generated in this work; see Methods and SI) revealed that W96 engages in a cation–*π* interaction with K52, as well as stabilizing hydrogen bonds with D101 and surrounding residues, including R94 (Figure 5a). This intricate network suggests that both the aromatic nature and precise orientation of W96 are critical for maintaining high-affinity binding. However, W96 is located near a negatively charged surface (Figure 5b), rendering it highly susceptible to oxidation. Under AAPH-induced oxidative stress, W96 exhibited a 97% oxidation rate, which led to a dramatic *>*500-fold reduction in binding affinity. A conventional W96F substitution resulted in complete loss of function, with a *>*1000-fold decrease in affinity, highlighting the limitations of such traditional mitigation strategies in this case.

To further evaluate the applicability of our model, we pursued a rational dual-optimization strategy aimed at reducing the oxidation risk of W96 while preserving binding. Notably, this optimization was performed without access to the CryoEM Fab–antigen complex at the time, highlighting the generalizability of our approach for early-stage discovery programs where fab-antigen complex structure is often unavailable.

The optimization was carried out over three iterative design rounds: 1) **Global charge tuning**, to increase overall isoelectric point (pI) and improve chemical stability. 2) **Local targeting**, where residues within 10 Å of W96 were mutated to modulate the local electrostatic potential. 3) **Combinatorial refinement**, based on binding and expression results from prior rounds.

Figure 5c shows that these charge-altering mutations were effective in raising the side-chain electrostatic potential (Epot) of W96, which correlated strongly with reduced oxidation rates. Figure 5d summarizes the top 13 variants, including Epot, oxidation reduction relative to the lead candidate, and their impact on binding (reported as fold change in K_D_). While some variants exhibited trade-offs between stability and affinity, two emerged as particularly promising.

Variant S11, featuring a single point mutation, achieved a 27.2% reduction in W96 oxidation with only a 1.4-fold reduction in binding affinity. Variant S13, with additional mutations, showed a 49.9% decrease in oxidation at the cost of a 20.7-fold reduction in affinity—still significantly better than the *>*1000-fold loss observed with W96F. These data demonstrate that it is possible to tune the local electrostatics around a critical Trp residue to reduce its oxidation risk, while retaining functional binding through careful, structure-guided design.

### Redox potential simulations reveal electrostatic control of Trp oxidation

To elucidate the mechanistic basis behind the differential oxidation propensity of Trp residues in distinct electrostatic environments, we performed redox potential molecular dynamics simulations using the redox replica exchange method. ^58^ Two antibody systems were selected for comparison: Mab4, which contains a Trp residue located in a region of negative electrostatic potential, and Mab4.RR, a variant in which two Asp residues are substituted with two Arg residues, resulting in a positively charged environment. This charge reversal leads to an 85% reduction in observed Trp oxidation in Mab4.RR, making it an ideal model system for probing the impact of local electrostatics on redox behavior.

In both systems, the Trp radical was modeled at the nitrogen atom of the indole ring. While radical formation is mechanistically more favored at the C2 or C3 carbon positions, ^36^ we chose the nitrogen site due to the availability of corresponding electrochemical data and associated free energy changes (Δ*G*) required to compute reduction potentials. ^59^ Our goal was not to pinpoint the exact radical species but to assess whether any Trp radical formation would exhibit a shift in redox potential under different electrostatic conditions.

Redox replica exchange MD simulations were conducted by titrating the Trp residue (TRX 100a) over a redox potential range of 900–1250 mV. The fraction of reduced species at each potential was used to determine the standard reduction potential (*E*^0^) of Trp in each system (see Figure 6). The results revealed that the Trp in Mab4 exhibited an *E*^0^ that was 88 mV lower than in Mab4.RR. Notably, the *E*^0^ of Mab4.RR closely matched that of a capped tryptophan control, indicating that the electrostatic environment in Mab4 leads to a significantly more oxidizable Trp residue. This 88 mV difference corresponds to a ∼1.5 pH unit shift in redox potential, underscoring the substantial influence of local electrostatics on Trp oxidation susceptibility.

**Figure 6:**
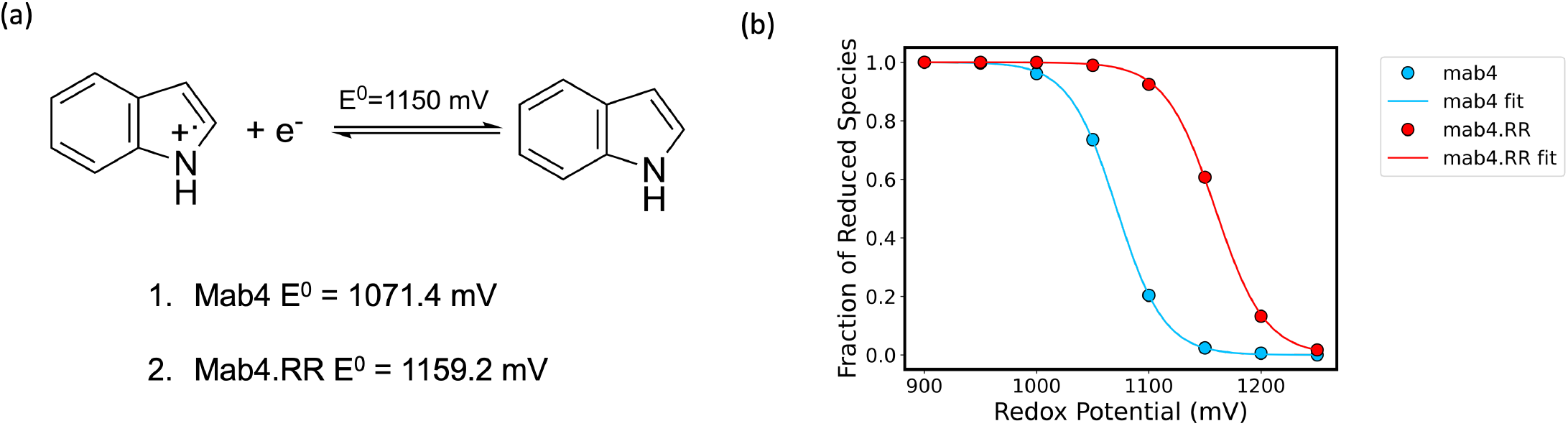
a) Calculated redox potential (E^0^) for Trp100a in Mab4 and its electrostatically engineered variant Mab4.RR. The lower E^0^ in Mab4 indicates higher susceptibility to oxidation, consistent with its negatively charged local environment. b) Redox titration curves for Trp100a in Mab4 (blue) and Mab4.RR (red), fitted using the Nernst equation. The upward shift in E^0^ in Mab4.RR reflects reduced oxidation propensity due to a more favorable (less negative) electrostatic potential surrounding the Trp residue.

These findings provide direct evidence that electrostatic modulation can significantly alter the redox potential of Trp residues in antibody frameworks, with potential implications for predicting and mitigating site-specific oxidation risks in therapeutic antibodies.

## Discussion

Oxidation remains a major degradation pathway in antibody therapeutics, potentially compromising binding affinity, stability, and overall developability. In this study, we analyzed experimentally measured oxidation rates for a diverse panel of 187 antibodies and developed machine learning classifiers to identify key structural determinants of Trp oxidation risk. Using expert-guided features and a combination of established and novel molecular descriptors, we confirmed that SASA is a primary driver of oxidation susceptibility. Importantly, we also identified local electrostatic potential (Epot) as a strong modulator of oxidation risk, with negative electrostatics significantly increasing the likelihood of oxidation.

A simplified two-parameter model incorporating just SASA and Epot achieved 79% classification accuracy, approaching the 84% performance of more complex, full-feature ML models. To validate the generalizability of these findings, we applied the model to a blind test set of eight clinical-stage antibodies containing 10 Trp residues in their CDRs. Despite substantial variation in sequence, target, and other molecular features, the model accurately predicted oxidation susceptibility, reinforcing the robustness of SASA and electrostatics as key descriptors.

The long-range nature of electrostatic effects supports a rational design strategy for mitigating oxidation through targeted point mutations. Introducing one to three charge-altering mutations near oxidation-prone Trp residues reduced oxidation rates by an average of 50% in four out of five antibodies tested, while maintaining binding affinity in two. Notably, several of these beneficial mutations were located in CDRs distinct from the one containing the oxidizing Trp, or at distant sites not in direct contact with the residue, highlighting the long-range effect of electrostatics on oxidation risk.

We further integrated these insights into a multi-parameter design strategy aimed at co-optimizing oxidation resistance and binding affinity in a clinical anti-CD33 antibody. This antibody contains a Trp96 residue that oxidizes at 97% upon AAPH stress. By shifting the Fv isoelectric point (pI) to more positive values and applying sequence-based affinity maturation, we generated several variants with substantially reduced oxidation while preserving antigen binding. The extent of binding loss in these variants was notably less than that observed in Trp-to-Phe substitutions, a common but often destabilizing engineering tactic.

To explore the mechanistic basis of electrostatic influence on oxidation, we conducted redox potential MD simulations on two closely related antibody variants. A pair of D→R mutations in Mab4 shifted the local electrostatic environment from negative to positive, and simulations revealed that the Trp in the positively charged context exhibited a significantly higher reduction potential (E^0^), consistent with a lower oxidation propensity. These simulations confirm that electrostatics alone can modulate the Gibbs free energy landscape for electron transfer, altering oxidation risk independently of other structural factors.

One mechanistic explanation for the role of electrostatics is that Trp oxidation involves electron loss, which is destabilized in positively charged environments due to electrostatic repulsion. This raises the energetic cost of oxidation while simultaneously disfavoring the reverse electron transfer. This is consistent with previous work showing that local electronic structure and polarity can redistribute electron density over the indole ring and facilitate reactions with singlet oxygen. ^60,61^ Further support comes from studies on azurin and other redox-active proteins, which show that local polarity and hydrogen bonding modulate Trp redox behavior. ^62–64^

While prior studies emphasized short-range water-mediated interactions and hydrogen bonding, our findings extend these principles by demonstrating the long-range impact of electrostatics, supported by Poisson-Boltzmann calculations and mutation experiments. Charge-altering mutations introduced at sites distant from a Trp residue significantly reduced its oxidation rate, broadening the scope of viable engineering strategies in antibody design.

One limitation of our study is its focus on AAPH-induced oxidation. Although other stressors, such as light, metal ions, or heat, also contribute to oxidation, ^65^ our redox MD simulations suggest that electrostatics influence oxidation independently of the stressor type by altering redox energetics. A potential AAPH-specific concern is its positive charge, which might preferentially localize to negatively charged regions and artifactually enhance oxidation. However, comparative experiments using a neutral AAPH analog on eight antibodies yielded nearly identical oxidation profiles (Figure S1), suggesting that AAPH functions primarily as a bulk-phase radical generator and does not require electrostatic attraction to exert its effect.

Thus, our electrostatic descriptors appear broadly applicable across different oxidation pathways, though future studies should experimentally test their relevance under diverse oxidative conditions.

In future work, structural data from Fab-antigen complexes could be incorporated to enable finer control over affinity- and stability-optimizing mutations. Such data would allow more precise tuning of electrostatics while preserving essential binding contacts. Nonetheless, even in the absence of structural models, early-stage sequence optimization can benefit from the multi-parameter strategy presented here.

In conclusion, we demonstrate that local side-chain electrostatics, in addition to the established solvent accessibility, is a key determinant of Trp oxidation risk in antibodies. We show that this feature is predictive, tunable, and mechanistically supported by redox potential simulations. Our findings offer a practical and generalizable framework for rational antibody design, enabling optimization of both oxidation stability and binding affinity, critical goals in therapeutic antibody development.

## Methods

### Antibody dataset and oxidation stress conditions

The internal dataset has about 187 unique mabs (light chain and heavy chain combination). These mabs were oxidized at pH 5.5 using 2-2’-azobis(2-amidinopropane) dihydrochloride (AAPH) for 16 hours at 37°C and the CDR Trp oxidation was reported with respect to a non-spiked control. A total of 264 CDR Trp residues were present in the 187 mabs, with 205 CDR Trp not oxidized and 59 data points being oxidized. A cutoff threshold of 35% oxidation rate was used for this classification. Similar to internal mabs, clinical stage antibodies and engineered mabs were oxidized using AAPH. Additionally, the clinical antibodies were oxidized using an AAPH variant (Figure S1 under same condition as AAPH oxidation stress study).

### Antibody fab structure modeling and molecular dynamics simulations

We constructed the variable domain (Fv) structure from the antibody sequences using the DeepAb^57^ modeling technique. To generate the full Fab structures, an internal VMD^66^ script was used to graft the Fvs onto the constant region of the Herceptin Fab by aligning their elbow loops. The decision to generate the Fab domain, rather than the Fv alone, was made to account for any potential electrostatic contributions from the Fab’s constant region. The resulting Fab structures were then subjected to molecular dynamics (MD) simulations. The Fab structure was solvated and parameterized using the FF14SB force field ^67^ and TIP3P^68^ water model in a 10 Å truncated octahedron box. Following relaxation protocol, 500 ns of MD simulation was performed. Further, 1000 snapshots were extracted every 0.5 ns for feature analysis. The full protocol is described in the SI.

### Feature extraction from MD trajectories

For each MD simulation frame, we extracted 50 features to describe the tryptophan oxidation propensity, as shown in Table S1. For each antibody in our dataset, we computed 50 residue-level structural features, capturing various characteristics such as solvent accessibility (e.g., SASA, SASA loop, n loop water 5), flexibility (e.g., loop rmsd, res rmsd), local hydrophobicity (e.g., Spatial Aggregation Propensity (SAP) scores), backbone dihedral angles (*ψ, ϕ*, and *χ* angles of neighboring residues), local electrostatic potentials (e.g., Epot), and interactions with neighboring polar, nonpolar, and aromatic residues. Detailed protocol is provided in the SI.

### Machine learning models and feature analysis

Feature relationships were assessed by hierarchical clustering. Dendrogram plot was generated using Spearman rank correlation matrix and single linkage clustering to visualize feature relationships. For quantifying feature importance to Trp oxidation rate, RF regressor (1000 estimators) was trained with standardized features using a 5-fold cross-validation split, with SHAP values computed on the test set using shap.TreeExplainer. Further GB and RF classifiers were used to predict oxidation rates as a binary classification problem. For classification, oxidation rates larger than 35% were used to create two classes (reactive vs non-reactive). Detailed ML protocol is explained in SI.

### Antigen binding kinetics

All surface plasmon resonance (SPR) measurements were conducted on a Biacore T200 instrument (Cytiva). Anti-human IgG antibody was immobilized on a CM5 chip according to the manufacturers recommendations. The mabs in question were then captured on the chip (1 µg/mL, 60 sec) and affinity for the respective antigen determined using single cycle kinetics. The sensograms were analyzed using the manufacturer’s software and fit to a 1:1 Langmuir binding model to calculate the kinetic and binding parameters. The full protocol is described in SI.

### Affinity determination of multi-property optimization clones using SPR

The binding affinity of protein and protein interaction was determined by SPR technology (Biacore™-8K+, Cytiva). Briefly, each antibody variant was captured by Protein A sensor chip (Series S) on the different flow cell to achieve approximately 150 response units (RU), followed by the injection of fivefold serial dilutions of human CD33 protein (R&D Systems; 0.16 nM to 100 nM) in HBS-EP buffer.

Sensorgrams were processed with reference and blank subtraction and analyzed using a 1:1 Langmuir binding model to calculate the kinetic and binding parameters. The detailed protocol is provided in SI.

### Multi-property optimization of affinity and tryptophan oxidation

Internal lead candidates were engineered in three iterative steps with distinct optimization objectives. Initially, a structurally-aware sequence optimization approach ^69^ was adapted to identify acidic framework residues (Asp/Glu) for substitution with basic residues (Arg/Lys) in tolerated positions, generating 24 candidate sequences for small-scale expression and affinity measurements. Next, mutations were introduced to Ala, Lys, and Arg around the oxidizing Trp residue to increase affinity, identifying single mutations, light chain double mutations, and combinations of heavy chain double and triple mutations. In the final step, beneficial mutations from both previous stages were combined in silico, producing 23,915 candidate sequences. Active learning ^70^ was used to select 14 sequences with desired properties, which were tested for improved affinity and oxidation resistance, demonstrating the effectiveness of multi-property optimization via active learning-assisted combinatorial mutagenesis. The detail protocol is provided in SI.

### Redox MD setup

The Mab4 WT and Mab4.RR variants were used for redox replica exchange simulations, with the tryptophan (TRX) residue parametrized as titratable in two states (Trp and Trp^.+^). Redox MD was performed on a capped tryptophan dipeptide, and the Trp 100a in Mab4 was defined as the titratable residue. Simulations included 8 redox windows from 900 mV to 1250 mV, with replica exchange runs of 60 ns, 10,000 exchange attempts, and solvent relaxation after each exchange. Full methods are provided in the SI.

### Cryogenic electron microscopy data collection, processing, and model refinement

Cryo-EM data was collected using a Titan Krios G3i with a K3 camera, capturing over 18,000 movies at varying tilts. The data were processed with cryoSPARC, including motion correction, CTF estimation, and 2D classification, resulting in a 3D reconstruction at 3.5 Å resolution. Model building was done using UCSF Chimera ^71^ and the human CD33 PDB structure, followed by refinement in COOT^72^ and Phenix. ^73^ The final model was validated and refined to 3.5 Å resolution. Full details of the method are provided in the SI.

## Author contributions

S.L. and S.I. conceived the project and designed the study. S.L. performed data curation, MD simulations, descriptor calculations, data analysis, variant designs, and validation. S.M. and N.J. measured binding affinities for Mabs 1–4 and their variants.N.S. conducted AAPH stress experiments and oxidation measurements. F.S., W.-C.L., and A.W. generated CD33 data and carried out multi-parameter optimization. R.K. and S.W. designed and analyzed Mab5 variant data. J.Z. contributed clinical antibody oxidation data and designed AAPH variant. R.P. and C.A. determined the CD33 Cryo-EM structure. F.J.I. supported descriptor calculations. C.C., M.H., and Y.W. engineered Mabs 1, 2, 4, and CD33. S.A. and B.K. contributed research concept and mechanistic insights; assisted variant designs. S.I. supervised the project, developed descriptors and machine learning models, performed feature analyses, and co-designed mutations. S.L. and S.I. wrote the manuscript with input from all co-authors.

## Acknowledgement

The authors thank Trevor Swartz, Tomasz Baginski, Nandhini Rajagopal for helpful comments on the manuscript.

## Declarations

All work described in this paper was funded by Genentech Inc., South San Francisco. All authors are or were employees of Genentech Inc. (a member of the Roche Group).

