## Supplementary Information for "Rational design of oxidation-resistant antibodies through local electrostatic modulation"

‖*Structural Biology, Genentech, South San Francisco, United States*

### Methods

#### Detailed molecular dynamics protocol

The fab protein was placed in an explicit solvent model using tleap. The protein was build using FF14SB,[1] and for water TIP3P[2] model was used. A truncated octahedron water box was used with the edge of the box being 10 Å from the protein. All simulations were performed on Amber 20.[3] The initial solvated fab structure was first minimized using 1000 steps steepest descent and 4000 steps conjugate gradient with a positional restraint on the protein backbone. The system was then heated gradually from 10 K to 300 K in the NVT ensemble, and additional NVT run for 2.3 ns at 300 K was conducted. During the heating phase, the protein is restrained by a force constant of 1.0 kcal $mol^{-1}$Å$^{-2}$. The timestep used in these calculations is 4 fs, achieved by HMass repartitioning.[4] Further, the system was equilibrated in an NPT ensemble for 8 ns using Monte carlo barostat and langevin dynamics with the protein restrained by a force constant of 1.0 kcal $mol^{-1}$Å$^{-2}$. The final round of relaxation consists of 10 ns run in an NVT ensemble with no restraints. After the relaxation, 500 ns of dynamics was conducted in the NVT ensemble. From this molecular dynamics (MD) simulation, 1000 snapshots (pdb files) corresponding to every 0.5 ns interval were extracted for the feature analysis.

#### Feature calculations

For each frame from MD simulations, we extracted 50 features to describe the propensity of tryptophan (Trp) oxidation, as shown in Table S1. These features can be divided into three broad categories- Trp neighboring residues, electrostatics, and hydrophobicity. The neighboring residues category further consists of backbone and sidechain angle ($\phi$, $\psi$, $\chi$) for the Trp, the angles for residues preceding and following tryptophan, root mean square deviation (RMSD) during the MD run of the Trp and loop (two residues preceding and following Trp, five residues in total), the number of waters and acidic residues oxygen and sulfur atoms within 3 and 5 Å of the Trp (n_res_water_3, n_quench_3), the solvent accessibility and exposure of the Trp, the number of polar, nonpolar, aromatic, acidic and basic residues. The electrostatics feature category consists of the APBS[5] electrostatics contribution around the Trp residue and the CDR loop where the Trp is present for the pdbs at pH 7. The broader hydrophobicity category consisted of the hydrophobicity and hydropathy scale for the Trp and residues within 3 and 5 Å of the Trp, the spatial aggregation propensity (SAP) of the Trp using different hydrophobicity scales such as Black and Mould (BM), Kyte-Doolittle (KD), Wimley-White (WW). The features related to the angle, neighboring residues, and SASA_res are evaluated by ptraj[6] and mdtraj.[7] The electrostatics Epot features were calculated using APBS[5]and the Moldesk tool.[8] The feature SASA and Exposure in Table S1 were evaluated with MSMS[9] tool.

Table S1: Features used to describe Trp oxidation for Machine Learning

| Feature | Description |
|---|---|
| Neighboring residues | |
| n_$\phi$ | $\phi$ dihedral angle of Trp |
| n-1_$\phi$ | $\phi$ angle of the residue before Trp |
| n-1_$\psi$ | $\psi$ angle of the residue before Trp |
| n-1_$\chi$ | $\chi$ angle of the residue before Trp |
| n+1_$\phi$ | $\phi$ angle of the residue after Trp |
| n+1_$\psi$ | $\psi$ angle of the residue after Trp |
| n+1_$\chi$ | $\chi$ angle of the residue after Trp |
| res_rmsd | rmsd of Trp |
| loop_rmsd | rmsd of total 7 residues including Trp (n-3 to n+3) |
| n_res_water_3 | # of water around 3Åof Trp |
| n_loop_water_3 | # of water around 3Åof Trp for n-3 to n+3 residues |
| n_quench_3 | # of OD1, OD2, OE1, OE2, SD, SG, OXT around 3 Åof Trp |
| n_res_water_5 | # of water around 5 Åof Trp |
| n_loop_water_5 | # of water around 5 Åof Trp for n-3 to n+3 residues |
| n_quench_5 | # of OD1, OD2, OE1, OE2, SD, SG, OXT around 5 Åof Trp |
| SASA_res | sasa of by shrake rupley Trp, probe radius 1.6 Å |
| SASA_Loop | sasa of by shrake rupley n-3 to n+3 residue, probe radius 1.6 Å |
| SASA | sasa of by MSMS Trp, probe radius 1.4 Å |
| Exposure | %sasa by MSMS keeping a cutoff for free Trp as 266.38 Å$^2$ |
| Polar_Residues_Hbond | # of polar residues that form hydrogen bond with Trp using cpptraj |
| Polar_Residues_Contact | # of polar residues that form native contacts with Trp using cpptraj |
| Nonpolar_Residues_Contact | # of nonpolar residues that form native contacts with Trp |
| Aromatic_Residues_Contact | # of aromatic residues that form native contacts with Trp |
| Acidic_Residues_Contact | # of acidic residues that form native contacts with Trp |
| Basic_Residues_Contact | # of basic residues that form native contacts with Trp |
| Electrostatic behavior | |
| Epot_Sum | electrostatic potential of Trp using APBS |
| Epot_Sum_Neg | negative electrostatic potential of Trp using APBS |
| Epot_Sum_Pos | positive electrostatic potential of Trp using APBS |
| Epot_loop_Sum | Electrostatic potential for the CDR loop where Trp is present |
| Epot_loop_Sum_Neg | Negative Electrostatic potential for the CDR loop where Trp is present |
| Epot_loop_Sum_Pos | Positive Electrostatic potential for the CDR loop where Trp is present |
| nb_sph_3_q | total Charge for neighbors around Trp at 3 Å |
| nb_sph_5_q | total Charge for neighbors around Trp at 5 Å |
| Hydrophobic behavior | |
| nb_seq_score_1 | Hydrophobicity score for 5 residues, n being Trp (n-2 to n+2) |
| nb_seq_score_2 | Hydropathy score for 5 residues, n being Trp (n-2to n+2) |
| nb_sph_3_score_1 | Hydrophobicity score for neighbors around 3 Åof Trp |
| nb_sph_3_score_2 | Hydropathy score for neighbors around 3 Åof Trp |
| nb_sph_5_score_1 | Hydrophobicity score for neighbors around 5 Åof Trp |
| nb_sph_5_score_2 | Hydropathy score for neighbors around 5 Åof Trp |
| SAP_BM_Sum | Spatial aggregation propensity of Trp using BM scale total sum |
| SAP_BM_Sum Neg | SAP of Trp using BM scale negative values sum |
| SAP_BM_Sum Pos | SAP of Trp using BM scale positive values sum |
| SAP_KD_Sum | SAP of Trp using KD scale total sum |
| SAP_KD_Sum Neg | SAP of Trp using KD scale negative values sum |
| SAP_KD_Sum Pos | SAP of Trp using KD scale positive values sum |
| SAP_WW_Sum | SAP of Trp using WW scale total sum |
| SAP_WW_Sum_Neg | SAP of Trp using WW scale negative values sum |
| SAP_WW_Sum_Pos | SAP of Trp using WW scale positive values sum |

#### Machine learning models and feature analysis

**Dendrogram plot:** The relationships between features were evaluated through hierarchical clustering. A Spearman rank correlation matrix was computed to assess monotonic associations among features. Single-linkage hierarchical clustering was then applied to the correlation matrix, producing a dendrogram that visually represents feature clustering based on their correlation distance.

**SHAP (SHapley Additive exPlanations) plot:** SHAP and Random Forest regression was used to identify features predictive of oxidation rates features that were standardized with StandardScaler to ensure uniform scale. A 5-fold cross-validation split was implemented with the KFold method, allowing one train-test split to be selected for SHAP analysis. A Random Forest regressor model with 1000 estimators was trained on the training split, with SHAP values computed on the test set using shap.TreeExplainer. Mean absolute SHAP values were calculated per feature to quantify feature importance. A SHAP summary plot was also generated.

**Gradient Boosting (GB) and Random Forest (RF) Classifiers:** GB and RF classifiers were used to predict oxidation rates as a binary classification problem. For classification, oxidation rates $> 35$ % were used to create two classes (reactive vs non-reactive). A GB classifier with 20,000 estimators and a RF classifier with 10,000 estimators were used, each trained using 10-fold cross-validation to ensure robust performance estimation. The performance metrics calculated included sensitivity, specificity, accuracy, and Matthews Correlation Coefficient (MCC), based on the confusion matrix derived from model predictions.

#### Antigen binding kinetics

All surface plasmon resonance (SPR) measurements were conducted on a Biacore T200 instrument (Cytiva). Anti-human IgG antibody was immobilized on a CM5 chip according to the manufacturers recommendations. The mabs in question were then captured on the chip (1 µg/mL, 60 sec) and affinity for the respective antigen determined using single cycle kinetics. 0.01 M HEPES pH 7.4, 0.15 M NaCl, 0.005% v/v Surfactant P20 was used as the running buffer. Antigen concentrations were varied to bracket the KD and dissociation was monitored for 600-1800 seconds after injection. The sensograms were corrected by subtracting from the blank flow cell as well as a zero point run (0 nM antigen). The corrected sensograms were analyzed using the manufacturer's software and fit to a 1:1 Langmuir binding model to calculate the kinetic and binding parameters.

#### Affinity determination of multi-property optimization clones using SPR

The binding affinity of protein and protein interaction was determined by SPR technology (Biacore™-8K+, Cytiva). Briefly, each antibody variant was captured by Protein A sensor chip (Series S) on the different flow cell to achieve approximately 150 response units (RU), followed by the injection of five-fold serial dilutions of human CD33 protein (R&D Systems; 0.16 nM to 100 nM) in HBS-EP buffer (100 mM 4-(2-hydroxyethyl)-1-piperazineethanesulfonic acid (HEPES) pH 7.4, 150 mM NaCl, 3 mM EDTA, 0.05% (v/v) Surfactant P20) with a standard flow rate of 50 µl/min at 25°C to reach a maximum binding response around 50 RU. The sensograms were recorded, processed by reference and blank subtraction, and evaluated by a simple one-to-one Langmuir binding model (Biacore™ Insight Software) to determine association rate constant ($k_{on}$), dissociation rate constant ($k_{off}$) and the equilibrium constant (KD).

#### Multi-property optimization of affinity and tryptophan oxidation

Variants of the internal lead candidates were engineered in 3 iterative steps with different optimization objectives. First, a structurally-aware sequence optimization approach[10] was adapted to manually identify acidic framework residues (Asp/Glu) suitable for substitution with basic residues (Arg/Lys) in tolerated positions (e.g., AHO HC: 84, 92; AHO LC: 17, 88, 97). This yielded 24 candidate sequences with edit distances of 1-4 Å, selected for automated small-scale expression, purification, and affinity measurements using SPR.[11]

Second, we explored substitutions to Ala, Lys, and Arg residues within a 10 Å radius of the oxidizing Trp. Mutations were screened to achieve ≤30-fold increases in affinity. This stage identified single mutations, light chain double mutations, and random combinations of heavy chain double and triple mutations, tested with their respective seed light or heavy chains.

Finally, beneficial mutations from Steps 1 and 2 were combined in silico, generating 23,915 candidate sequences with edit distances of 2-11 Å across both chains. Active learning techniques[12] were used to select 90 sequences with desired properties: pKD > 6, calculated pI > 8.2, and a positive local Trp potential (Epot_POS). These optimized variants were tested for improved affinity and oxidation resistance.

#### Redox MD setup

The Mab4 WT and Mab4.RR were selected to perform redox replica exchange simulation. For performing redox MD, the tryptophan (TRX) residue was parametrized to be titratable. Two states for Trp were defined as State 1 (Trp) and State 2 ($Trp^{\cdot+}$). The charges for each atom were obtained using Gaussian 09 program[13] with HF/6-31G* level capped Trp dipeptide (ACE-TRX-NME) as shown in Table S2. Constant redox potential MD was performed on capped Trp dipeptide (ACE-TRX-NME) and using Metropolis Monte Carlo redox state change attempt similar to previous publications,[14] the $\Delta G_{elec,ref}$ term was fitted to predict $E^o$. Next the TRX 100a in Mab4 was defined as titratable tryptophan residue (TRX) and redox runs were conducted. Similar to previous MD setup minimization, heating and equilibration was done for the Mab4 and Mab4.RR with the titratable TRX residue. The last equilibration coordinates will be used as the starting points for all redox windows. A total

Table S2: Charge distribution for the Redox Active Tryptophan (TRX) Residue

| | State 1 (Trp) | State 2 ($Trp^{\cdot+}$) |
|---|---|---|
| Atom | Charge (a.u.) | |
| N | -0.4157 | -0.4157 |
| H | 0.2719 | 0.2719 |
| CA | -0.0014 | -0.0014 |
| HA | 0.0876 | 0.0876 |
| CB | -0.019 | 0.1487 |
| HB2 | 0.083 | 0.098 |
| HB3 | 0.083 | 0.098 |
| CG | -0.3931 | -0.6591 |
| CD1 | -0.0479 | 0.1923 |
| HD1 | 0.1891 | 0.09 |
| NE1 | -0.5539 | -0.843 |
| HE1 | 0.3844 | 0 |
| CE2 | 0.2989 | 0.5922 |
| CZ2 | -0.3864 | -0.392 |
| HZ2 | 0.1842 | 0.1343 |
| CH2 | -0.1702 | -0.2215 |
| HH2 | 0.1495 | 0.0997 |
| CZ3 | -0.2037 | -0.2743 |
| HZ3 | 0.1594 | 0.1004 |
| CE3 | -0.3814 | -0.4736 |
| HE3 | 0.2675 | 0.2376 |
| CD2 | 0.3218 | 0.2769 |
| C | 0.5973 | 0.5973 |
| O | -0.5679 | -0.5679 |

of 8 windows ranging for 900 mV to 1250 mV in intervals of 50 mV, were used for redox run. For each potential, an initial equilibration of 20 ns with a redox change attempt every 500 steps at constant volume were done. These equilibrated structures at each potential were used for the redox replica exchange run. Two replicates of 60 ns redox replica exchange run were conducted with 10,000 number of times exchange between redox potentials was attempted and 10 ps of solvent relaxation after every exchange. The first 10 ns was considered as equilibration and the last 50 ns was used to evaluate the redox potential using cestats.[14]

#### Fab-antigen complex reconstitution

Human CD33 (residues 17-260) with N-terminal signal peptide sequence (MGGTAARLGAVILFVVIVGLHGVRG) and C-terminal Avi tag and TEV-cleavable Flag tag was expressed

Table S3: Cryo-EM data collection and refinement statistics

| Data collection and image processing | |
| --- | --- |
| Microscope | ThermoFisher Titan Krios |
| Detector | Gatan K2 |
| Voltage (kV) | 300 |
| Magnification | 105,000x |
| Defocus range (µm) | -0.5 to -2.5 |
| Pixel size (Å) | 0.838 |
| Electron exposure ($e^-$/Å$^2$) | 64 |
| Images (number) | 13,447 |
| Initial particles (number) | 7,725,681 |
| Final particles (number) | 561,118 |
| Symmetry imposed | C1 |
| Map sharpening, B factor (Å$^2$) | 127 |
| Map resolution (Å) | 3.5 |
| FSC threshold | 0.143 |
| Refinement | |
| Initial models used | PDB ID: 5IHB, MOE |
| Model resolution (Å) | 3.6 |
| FSC threshold | 0.5 |
| Model composition | |
| Chains | 3 |
| Non-hydrogen atoms | 6636 |
| Protein residues | 533 |
| Ligands | 0 |
| Mean B factors (Å$^2$) | |
| Protein | 54.02 |
| R.m.s. deviations | |
| Bond lengths (Å) | 0.003 |
| Bond angles (°) | 0.453 |
| Validation | |
| MolProbity score | 1.83 |
| Clash score | 14.54 |
| Rotamer outliers | 0.41 |
| Ramachandran plot | |
| Favored (%) | 97.06 |
| Allowed (%) | 2.94 |
| Disallowed (%) | 0 |
| Rama-Z | |
| Whole (N = 548) | -0.15 |
| Helix (N = 15) | 1.96 |
| Sheet (N = 250) | -0.26 |
| Loop (N = 283) | 0.11 |

in Human Embryonic Kidney (HEK) 293 cells in the presence of 1 mg/ml kifunensine. Six days post-transfection, the supernatant was collected and passed through an Anti-Flag

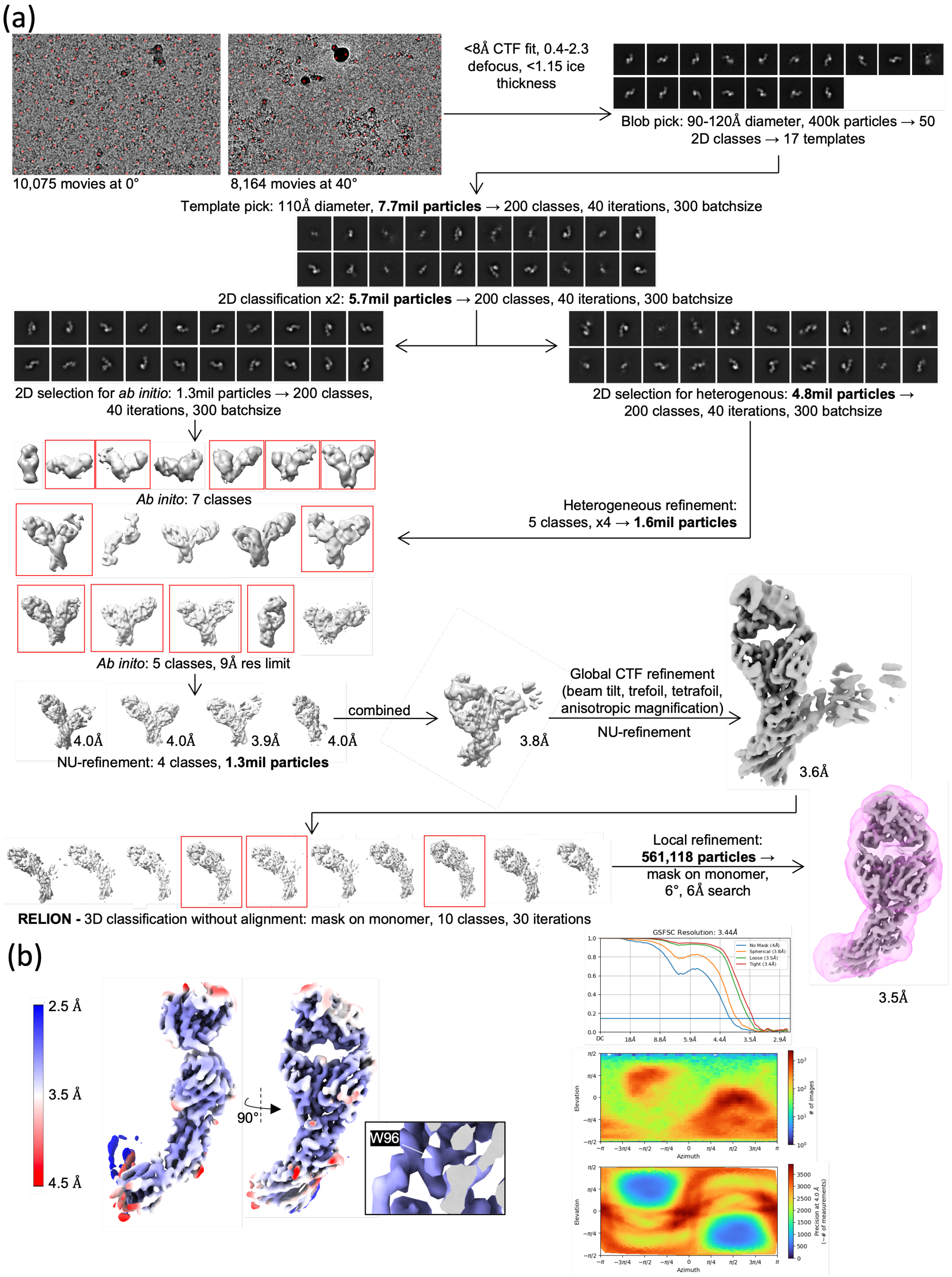

Figure S1: a) Processing workflow for fab antigen complex. All processing in cryoSPARC unless noted. Final FSC and view distribution plots are from local refinement. b) Local resolution map for Fab-antigen complex (local refinement map) with inset zoom on W96.

resin. Bound proteins were eluted using 0.1 M glycine pH 2.7-3.0 and further purified using Superdex S200 pg (Cytiva) equilibrated with 20 mM HEPES pH 7.5 and 150 mM NaCl. Fractions containing recombinant human CD3317-260-Avi-Flag protein were pooled, concentrated and snapped frozen using liquid nitrogen before storing at -80 °C. Fab-antigen complex was prepared by mixing purified Fab generated from Lys C digestion of anti-CD33 mAb and recombinant human CD3317-260-Avi-Flag deglycosylated with endo H and NEB protein deglycosylation mix II kit at 1:1 ratio followed by size exclusion chromatography using Superdex S200 pg (Cytiva).

Table S4: Sequences for the clinical antibodies

| Mab | $V_H$ Sequence | $V_L$Sequence |
|---|---|---|
| MabA | QMQLVESGGGVVQPGRSLRLSCAASGFT<br>FRTYGMHWVRQAPGKGLEWVAVIWYD<br>GSNKHYADSVKGRFTITRDNSKNTLNLQ<br>MNSLRAEDTAVYYCARAPQWELVHEAF<br>DIWGQGTMVTVSS | SYVLTQPPSVSVAPGQTARITCGGNNLGSK<br>SVHWYQQKPGQAPVLVVYDDSDRPSWIPE<br>RFSGSNSGNTATLTISRGEAGDEADYYCQV<br>WDSSSDHVVFGGGTKLTVL |
| MabB | QVQLVESGGGVVQPGRSLRLSCAASGFT<br>FFSYAMHWVRQTPGKGLEWVAVIWFDG<br>SNENYVDSVKGRFTISRDNSKNTLYLQMN<br>TLRAEDTAVYYCARDAWSYFDYWGQGTL<br>VTVSS | VIQLTQSPSSLSASVGDRVTITCRASQGISRA<br>LAWYQQKPGKGPKLLIYDASSLESGVPSRF<br>SGSGSGTDFTLTISSLQPEDFATYYCQQFNS<br>YPLTFGGGTKVEIK |
| MabC | EVQLVESGGGLVQPGGSLRLSCAASGFT<br>LSGDWIHWVRQAPGKGLEWVGEISAAG<br>GYTDYADSVKGRFTISADTSKNTAYLQM<br>NSLRAEDTAVYYCARESRVSFEAAMDYW<br>GQGTLVTVSS | DIQMTQSPSSLSASVGDRVTITCRASQNIAT<br>DVAWYQQKPGKAPKLLIYSASFLYSGVPSR<br>FSGSGSGTDFTLTISSLQPEDFATYYCQQSE<br>PEPYTFGQGTKVEIK |
| MabD | QVQLVQSGAEVKKPGASVKVSCKASGYI<br>FSNYWIQWVRQAPGQGLEWMGEILPGS<br>GSTEYTENFKDRVTMTRDTSTSTVYMEL<br>SSLRSEDTAVYYCARYFFGSSPNWYFDVW<br>GQGTLVTVSS | DIQMTQSPSSLSASVGDRVTITCGASENIYG<br>ALNWYQQKPGKAPKLLIYGATNLADGVPSR<br>FSGSGSGTDFTLTISSLQPEDFATYYCQNVLN<br>TPLTFGQGTKVEIK |
| MabE | QVQLVESGGGVVQPGRSLRLSCAASGFK<br>FSGYGMHWVRQAPGKGLEWVAVIWYDG<br>SKKYYVDSVKGRFTISRDNSKNTLYLQMN<br>SLRAEDTAVYYCARQMGYWHFDLWGRG<br>TLVTVSS | EIVLTQSPATLSLSPGERATLSCRASQSVSSYL<br>AWYQQKPGQAPRLLIYDASNRATGIPARFSG<br>SGSGTDFTLTISSLEPEDFAVYYCQQRSNWPP<br>LTFGGGTKVEIK |
| MabF | QVQLQQSGAELARPGASVKMSCKASGYT<br>FTSYTMHWVKQRPGQGLEWIGYINPSSG<br>YTNYIQRFKDKATLTADKSSSTAYMQVSSL<br>TSEDSAVYYCARGSRYDYYGMDYWGQG<br>TSVTVSS | QIVLTQSPAIMSASPGEKVTMTCSASSSVSYMH<br>WYQQKSGTSPKRWIYDTSKLASGVPARFSGS<br>GSGTSYSLTISSMEAEDAATYYCQQWSS<br>NPLTFGAGTKLELK |
| MabG | EVQLVESGGGLVQPGGSLRLSCAASGFTFS<br>SYGMDWVRQAPGKGLEWVSSIRGSRGSTY<br>YADSVKGRFTISRDNSKNTLYLQMNSLRAE<br>DTAVYYCARLYRYWFDYWGQGTLVTVSS | SYELTQPPSVSVSPGQTARITCSGDNLPKYYAH<br>WYQQKPGQAPVVVIFYDVNRPSGIPERFSGSNS<br>GNTATLTISGTQAMDEADYYCQAWWSSTPVFG<br>GGTKLTVL |
| MabH | QVQLVQSGAEVKKPGASVKISCKVSGYTLR<br>GYWIEWVRQAPGKGLEWIGQILPGTGRTN<br>YNEKFKGRVTMTADTSTDTAYMELSSLRSE<br>DTAVYYCARFDGNYGYYAMDYWGQGTTV<br>TVSS | QAVVTQEPSLTVSPGGTVTLTCRSSTGAVTTSN<br>YANWFQQKPGQAPRTLIGGTNNRAPGVPARFS<br>GSLLGGKAALTLSGAQPEDEAEYYCALWYSNH<br>WVFGGGTKLTVL |
| | Heavy chain | Light chain |
| Constant region sequence | ASTKGPSVFPLAPSSKSTSGGTAALGCLVKD<br>YFPEPVTVSWNSGALTSGVHTFPAVLQSSGL<br>YSLSSVVTVPSSSLGTQTYICNVNHKPSNTKV<br>DKKVEPKSCDKTHTCPPCPAPELLGGPSVFL<br>FPPKPKDTLMISRTPEVTCVVVDVSHEDPEV<br>KFNWYVDGVEVHNAKTKPREEQYNSTYRV<br>VSVLTVLHQDWLNGKEYKCKVSNKALPAPI<br>EKTISKAKGQPREPQVYTLPPSREEMTKNQ<br>VSLTCLVKGFYPSDIAVEWESNGQPENNYKT<br>TPPVLDSDGSFFLYSKLTVDKSRWQQGNVFS<br>CSVMHEALHNHYTQKSLSLSPGK | RTVAAPSVFIFPPSDEQLKSGTASVVCLLNNFYP<br>REAKVQWKVDNALQSGNSQESVTEQDSKDSTY<br>SLSSTLTLSKADYEKHKVYACEVTHQGLSSPVTK<br>SFNRGEC |

#### Cryogenic electron microscopy data collection and image processing

39 µl of the fab-antigen complex and 1 µl 40 mM BS3 was mixed and incubated for 10 min at 22°C. Reaction was quenched by adding 10 µl 1 M Tris pH 8.0 to the mixture. The sample was diluted to 0.25 mg/ml in 20 mM HEPES pH 7.5, 150 mM NaCl directly before vitrification, and 3 µl sample was applied to glow-discharged holey gold 0.6/1 300-mesh grids (Quantifoil). Grids were blotted for 5 s at 0 force before being plunge vitrified in liquid ethane using a MarkIV Vitrobot (ThermoFisher). The blotting chamber was maintained at 4 °C and 100% humidity during freezing.

Movies were collected using a Titan Krios G3i (ThermoFisher) outfitted with a K3 camera and Bioquantum energy filter (Gatan). The K3 detector was operated in nonCDS counting mode and the energy filter slit width was set to 20 eV. Movies were collected at a nominal magnification of 105,000×, physical pixel size 0.838 Å/pixel, with a 50 µm C2 aperture and 100 µm objective aperture at a dose rate of 15$e^-$/pixel.s. A total dose of 64 $e^-/Å^2$ was collected as a 59-frame movie, resulting in a 3 s movie with 1.08 $e^-$ per frame. Data were collected using semi-automated imaging scripts in SerialEM. 10,075 movies were collected in a 3 x 3 image shift pattern at 0° tilt and 8,164 were collected at 40° tilt. Beam tilt was corrected for using SerialEM's image shift vs coma calibration.

Data were processed on-the-fly through 2D classification using cryoSPARC v4.3.0. Movies were motion corrected using patch motion correction, CTF information was estimated using patch CTF estimation, and micrographs were curated based off of CTF resolution fit, defocus range, and ice thickness. Templates chosen from an initial blob pick were used for particle picking. Template picking yielded 7,725,681 picks. These were extracted in a 420-pixel box, and binned to 64 pixels. Two rounds of 2D classification were run with 200 classes, 40 online iterations, and 300 particle batch size. Seven ab initio classes were generated from 1,317,237 particles. Class averages with nominal particle shape were selected and 4,838,166 particles were moved into 3D sorting. These particles were subjected to four rounds of heterogeneous refinement using five of the seven ab initio volumes as starting models. Repetitive models were removed. 1,645,724 particles were re-extracted in a 450-pixel box, and binned to 256 pixels. This final pixel size was 1.3748 Å/pixel. A new ab initio reconstruction job was run followed by a heterogeneous refinement job with all of five initial models. Four of these classes were run through non-uniform refinement and yielded reasonable reconstructions of a monomer bound to Fab, with the second monomer at varying levels of resolution. All particles were combined into a single NU-refinement job to determine initial refinement angles. After global CTF refinement of beam tilt, trefoil, spherical aberration, tetrafoil, and anisotropic magnification, a consensus refinement of 1,380,263 particles refined to 3.5 Å. These particles were exported to RELION[15] and classified in 3D without refinement with a mask around a single monomer and Fab. Three classes showed the most contiguous density for the antigen and Fab. These 561,118 particles were reimported into cryoSPARC and local refinements were run with a 6° and 6 Å search with a mask around the full monomer-Fab, the IgV domain-Fab, and the IgV domain-Fv. The final maps was estimated to be 3.5, 3.4, and 3.4 Å by gold-standard FSC, respectively. Local resolution, orientation diagnostics, and local filtering jobs were run on the final map.

#### Model building and refinement

Initial models for rigid-body fitting into EM map using UCSF Chimera[16] were obtained using MOE[17] Fab structure prediction and available PDB crystal structure of human CD33 (PDB ID: 5IHB). The fitted model was manually inspected and rebuilt into EM densities using COOT version 0.9.8.93,[18] refined using real space refinement in Phenix version 1.21rc1-5015,[19] and validated using Molprobity.[20] Params for EM data collection and refinement statistics are presented in Table S3.

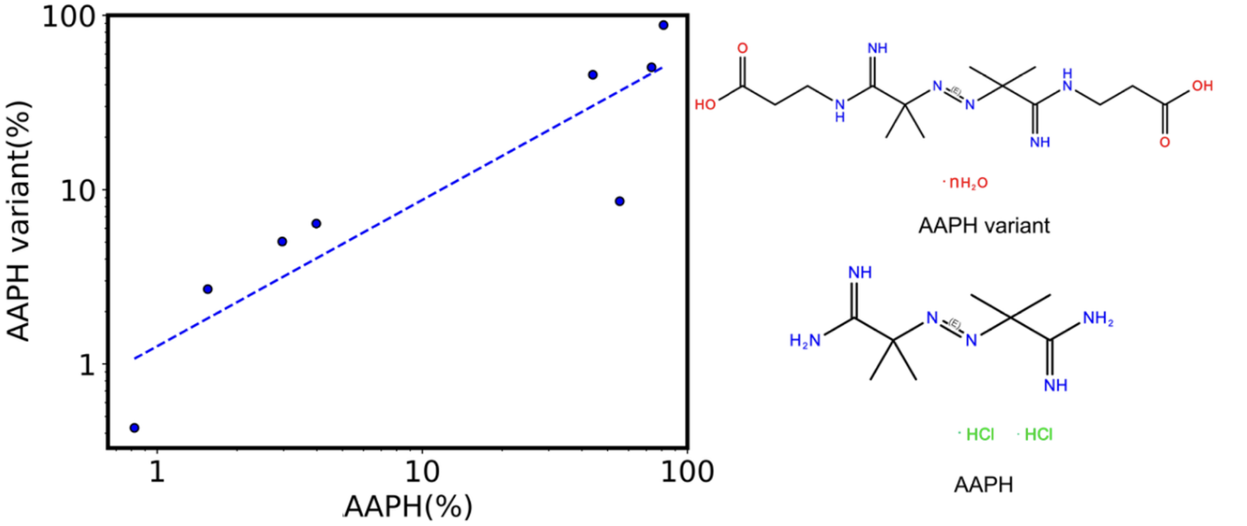

Figure S2: Impact of AAPH variant on measured oxidation rates of CDR Trp residues across eight clinical antibodies. Oxidation rates were assessed under stress conditions using both AAPH and its uncharged variant to compare their effects.
